# Fast and Slow Gene Expression Changes in Blood Following Acute Social Stress

**DOI:** 10.64898/2026.01.30.702738

**Authors:** Hampus Grönvall, Sylvia Abdelhalim, Fara Tabrizi, Sini Ezer, Gamze Yazgeldi Gunaydin, Erik Arner, Shintaro Katayama, Juha Kere, Fredrik Åhs, Lea Mikkola

## Abstract

Social stress is a risk factor for psychiatric disorders and also influences immune function. While it is known that acute social stress impacts the number of immune cells in circulation, the temporal dynamics of stress induced immune-related transcriptional changes in human blood remain unclear. To investigate changes in gene expression, we exposed 26 adults to the Trier Social Stress Test (TSST), and collected blood at baseline, as well as 5, 30, 60 and 90 min after stress. Whole-blood gene expression was profiled using a 5’ targeted RNA-sequencing method (STRT). Differential expression was analyzed using linear and cubic models. We observed a total of 54 differentially expressed genes following stress. Fast responses, with a transient peak immediately following stress, were enriched for cytotoxic T cell, NK cell and dendritic cell functions (e.g., *GZMB, GNLY, CCL4* and *GZMA*) and paralleled lymphocyte count changes. In contrast, gradual, linear responses without any evident peak were enriched for neutrophil related genes (e.g., *FPR2, PLAUR, CXCR2, AQP9*, and *QPCT*) and did not mirror neutrophil counts, indicating cell intrinsic transcriptional changes. From pathway and transcription factor enrichment analyses, IL-12 family mediated signaling is inferred as a central mechanism linking stress to immune gene regulation. Our results show that acute psychosocial stress induces both fast and slower changes in gene expression in different immune cell populations. The involvement of the IL-12-STAT4 axis and genes such as *PLAUR* and *FPR2* suggests molecular mechanisms through which stress-related immune activation may contribute to vulnerability for anxiety and depressive disorders.

## Introduction

Stress related psychiatric disorders impose a heavy burden on both individuals and society. An understand of the biological pathways linking stress to risk for psychiatric disorders is needed for finding new ways to inform prevention and intervention efforts. Chronic stress can influence the immune system, potentially increasing risk for depression and anxiety via inflammatory mechanisms[1]. Higher circulating levels of immune markers such as IL-6 and C-reactive protein have been associated with depression[2–4]. It has therefore been hypothesized that acute stress triggers gene expression changes in immune cells, and that repeated or prolonged activation of these immune pathways may contribute to the development of depression or anxiety after chronic stress exposure.

Prior studies that have used targeted approaches to study gene expression have shown that acute psychosocial stress can increase the expression of immune-related genes, such as the transcription factor NF-κB and cytokines IL-1β, IL-6 and IL10[5, 6]. While these findings point to stress induced activation of inflammatory signaling, unbiased whole-transcriptome studies are needed to discover novel stress responsive molecular pathways. To our knowledge, four genome wide transcriptomic studies of acute stress have been reported^7–10^. These studied all employed the Trier Social Stress Test (TSST)[11], but differed in sample characteristics (e.g., healthy volunteers vs. individuals with trauma history) and post-stress sampling times, ranging from immediately after stress to 3 h later. Furthermore, the specific genes showing the largest changes have been inconsistent between studies.

A major limitation of previous studies is the lack of concurrent immune cell count measurements alongside gene expression profiling. Acute stress is well known to cause rapid increases in circulating neutrophils, lymphocytes and monocytes, with neutrophils remaining elevated for an h or more following stress[12–14]. Such shifts in blood cell composition can confound bulk gene expression results. For example, when MacCormack et al.[15] adjusted for estimated cell-type proportion, many stress related gene expression differences became non-significant, suggesting that those signals were driven by changes in cell counts rather than gene regulation. Another gap in prior research is limited temporal resolution. Most studies collected at most one or two post stress samples, making it difficult to distinguish early gene responses from late responses.

To this end, we characterized whole genome changes in gene expression following stress by sequencing RNA isolated from whole blood at baseline, and at 5, 30, 60 and 90 min after stress. To compare changes in gene expression with changes in blood composition, we also measured immune cell counts at the same timepoints. This design allowed us to investigate which genes changed in activity, both early and late after stress.

## Subjects and Methods

### Participants

Participants were recruited from the Swedish Twin Registry[16] as part of the larger TwinStress research project. The study sample consisted of 26 participants (15 female, 11 male), between 32 and 58 years of age (M = 42, SD = 7.3). All participants signed informed consent, and the study was approved by the Swedish Ethical Review Authority (Dnr. 2020-00885).

### Procedure

To account for diurnal variation in cortisol levels, all sessions were conducted at approximately the same time of day, starting between 11:00 to 12:00. Upon arrival, a catheter was inserted in the non-dominant arm for blood sampling and participants were given a brief rest period before undergoing the Trier Social Stress Test (TSST)[11], a standardized psychosocial stress protocol. The TSST consisted of a simulated job interview: participants were given 3 min to mentally prepare a speech, then asked to deliver a 5 min oral presentation followed by a 5-min mental arithmetic task (counting backward from 2322 in steps of 13) in front of a two-person panel. A video camera was visible present to enhance social-evaluative threat.

### Blood sampling

Venous blood was collected at five timepoints: approximately 30 min prior to the TSST (baseline), and then at 5, 30, 60 and 90 min following the completion of the TSST. For the first three timepoints, plasma was isolated and analyzed for cortisol levels through electrochemiluminescence immunoassay (ECLIA) using a Roche Cobas e801 analyzer (Roche Diagnostics, Mannheim, Germany). Whole blood and plasma samples were stored refrigerated at 4 °C for a maximum of 24 h before being analyzed. Complete blood counts were determined using a Sysmex XN-series hematology analyzer (Sysmex Corporation, Kobe, Japan). These analyses were performed by the Karolinska University Laboratory at Karolinska Institutet, Stockholm.

### Statistical Analyses of Cortisol Levels and Cell Counts

To investigate changes across time in cortisol levels and blood cell counts, we fitted linear mixed-effects models using the lmerTest version 3.1.3[17] in R version 4.5.1[18]. Each model included timepoint as a fixed factor and participant ID as a random intercept to account for within-person dependencies. Tests of fixed effects were based on Type III F-tests with Satterthwaite-approximated degrees of freedom. Post-hoc contrasts between consecutive timepoints were estimated using emmeans version 2.0.1[19], with *p*-values adjusted for multiple comparisons via the Benjamini-Hochberg procedure within each outcome.

### Preprocessing, Cleaning and Quality assessment of STRT Sequencing Libraries

Whole-blood gene expression was profiled using single-cell tagged reverse transcription sequencing (STRT-Seq). Raw sequencing reads were processed with the STRT-N pipeline^20^. Sequencing reads were mapped to the reference human genome (hg38) and annotated using the GENCODE v43 basic annotation set^21^ via the UCSC Genome Browser^22^. We computed standard library-specific quality control (QC) metrics for each of the four multiplexed sequencing libraries (each containing 48 samples), including total mapped reads, spike-in reads, and 5’-end alignment rates, overall mapping ratio, ratio of mapped reads to spike-ins, and 5’-end coding read percentage. Based on these metrics outlier samples, non-template controls (NTCs), and technical duplicates were excluded from both libraries (see **Supplementary Figure S1**). Following quality control, 42, 34, 37 and 40 samples were retained for downstream analyses from each of the four libraries respectively.

Gene-level count matrices were generated after read alignment. Genes with zero counts in > 5 samples were filtered out to eliminate very low-expressed transcripts. Initially, 19 761 genes were detected, and after filtering 12 700 genes remained. Because our samples were sequenced in multiple library batches, we applied a method to correct for potential library-specific biases. Specifically, we used the Negative Binomial Generalized Linear Model – Library Bias Correction (NBGLM-LBC) package^23^ in R version 4.2.2[18] to adjust gene counts between libraries. Read depth information was extracted from BAM files using samtools^24^. We next performed spike-in-based variability analysis to identify genes with significant fluctuations across samples^25^. A total of 1171 genes showed high variability (Benjamini-Hochberg FDR corrected *p* < .05 in the spike-in variability test) and were labelled as fluctuating genes to be used in downstream analyses. Principal component analysis (PCA) was carried out on both raw and bias corrected count data using the Seurat package version 5.0.1^26^ to verify that library batch effects were effectively mitigated after correction (see **Supplementary Figure S2**). Following QC and batch correction, gene counts were spike-in normalized with log-transformation.

### Differential Gene Expression Analysis

Differential gene expression (DGE) analysis was carried out on the filtered and bias-corrected dataset of fluctuating genes using the R packages edgeR version 4.2.2^27^ and limma version 3.60.6^28^. We applied trimmed mean of M-values (TMM) normalization to account for library size differences. To capture potential non-linear trends in gene expression, we fit two separate regression models: a linear model and cubic polynomial model. In the linear model, expression was modelled as a function of time (in min) relative to TSST completion, treated as a continuous variable. In the cubic model, time was modeled with a third-degree (cubic) polynomial to allow for transient expression changes that peak and subside during the study window. Both models were applied to log2 counts-per-million values. Because our sample included for twin pairs, we accounted for within-pair non-independence using limmas duplicateCorrelation function. As sex and age were accounted for by the twin pair blocking, no other covariates were included in the models. From the resulting lmFit models, gene expression statistics were computed using limma’s eBayes function. Genes with Benjamini-Hochberg FDR corrected *p*-value < .05 in either model were deemed significantly differentially expressed.

### Gene Set Enrichment, Transcription Factor Enrichment, and Protein-Protein Interaction Analyses

To investigate enriched terms and pathways, we performed gene set enrichment analysis (GSEA), transcription factor enrichment analysis (TFEA), and protein-protein interaction analysis on the combined set of significant DE genes from both linear and cubic models (removing duplicates).

GSEA was performed using the g:Profiler web tool^29^ version e113_eg59_p19_6be52918 with g:OSt and default parameters. The results were then visualized using the multienrichjam ^30^ version 0.0.86.900, enrichplot version 1.28.1^31^and ggplot2 version 4.0.1^32^ packages alongside base R plotting functions. For TFEA, the ChEA3[33] transcription factor enrichment tool was used. The relationship between TFs and target genes from ChEA3 were then identified by clustering and visualization using using Clustergrammer[34]. Finally, STRING version 12.0[35] was used to investigate protein-protein interaction predictions. A full interaction network was generated using a medium confidence score cutoff (interaction score ≥ 400).

## Results

### Blood Cortisol Levels and Cell Counts

Cortisol varied significantly over time (*F*_2,49_ *=* 20.71, *p* < .001, 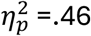, **Fig. 1A**). Consistent with the expected physiological stress response, there was an increase from baseline (M = 262.42, SD = 84.96) to 5 min after stress (M = 378.39, SD = 129.87, ΔM = 115.96, *p* < .001), followed by a decrease at 30 min after stress (M = 323.76, SD = 124.45, ΔM = -53.25, *p* = .005).

**Fig. 1.**
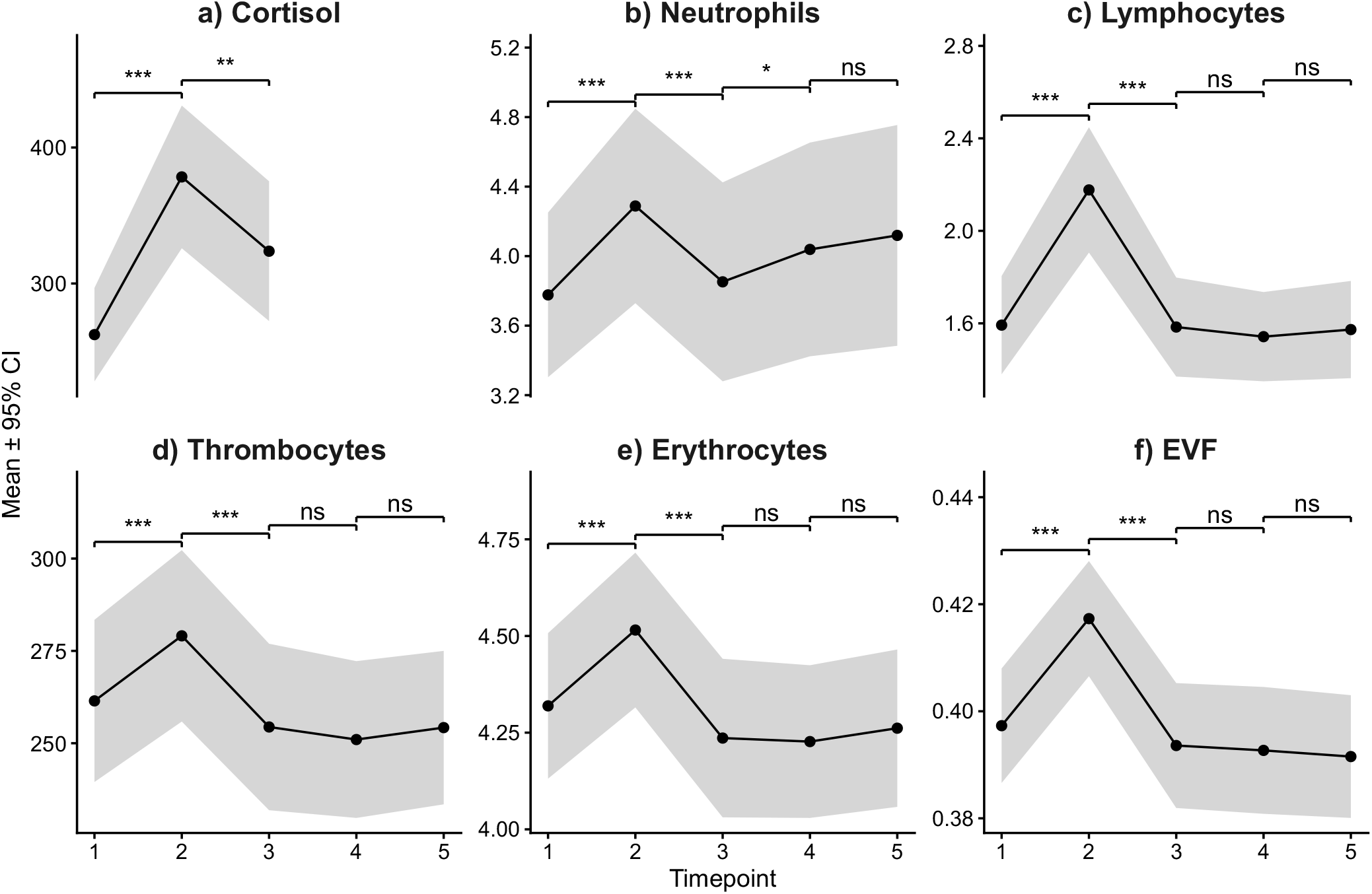
Cortisol Levels and Immune Cell Counts. Mean blood cortisol (nmol/L) and immune cell counts (10^9^/L) with shaded area representing 95 % confidence intervals at each timepoint. Significance levels for consecutive contrasts between timepoints are labelled: * *p* < .05, ** *p* < .01, *** *p* < .001, *ns* = non-significant.

As shown in **Fig. 1B-E**, all cell counts increased immediately after stress, before returning toward baseline. Neutrophil counts then increased again from 30 to 60 min, while other measures remained unchanged. Observed means, fixed effect estimates and contrasts across all timepoints available in **Supplementary Table S1-3**.

### Fast and Slow Gene Expression Changes After Acute Stress

We identified a total of 54 genes that were differentially expressed (DE) over time following the TSST (FDR adjusted *p-*value < .05 in at least one model; see **Table 1** for the top differentially expressed genes and **Supplementary Table S4-5** for the full results).

**Table 1.** Top Differently Expressed Genes. The five highest ranking genes by Log_2_ FC in either direction uniquely detected by the linear or cubic models, as well as the four genes that were found in both models. For linear models, Log_2_ FC represents change per min, while for polynomial models Log2FC represents spline components and do not have a straightforward biological interpretation. For this reason, Log_2_FC and p-values from the linear models are reported for the genes found in both models. Fast change genes indicate genes with a transient spike at 5 min post stress, detected b cubic model. Slow change genes indicate genes showing gradual change over 90 min without any obvious spike, detected by the linear model. Single Cell Type Cluster labels from The Human Protein Atlas[36]. Reported pvalues are Benjamini-Hochberg FDR adjusted.

| Gene | Single Cell Type Cluster | Change | Model | Log<sub>2</sub>FC | p |
| --- | --- | --- | --- | --- | --- |
| GNLY | NK-cells & T-cells - Adaptive immunity: Cytotoxicity | Fast | Both | -0.005 | .014 |
| GZMB | Plasmacytoid dendritic cells - Viral sensing and interferon response | Fast | Both | -0.004 | .029 |
| CCL4 | NK-cells & T-cells - Adaptive immunity: Cytotoxicity | Fast | Both | -0.004 | .026 |
| GZMA | NK-cells & T-cells - Adaptive immunity: Cytotoxicity | Fast | Both | -0.003 | .014 |
| PILRB | Non-specific - Mixed function | Fast | Cubic | 2.178 | .029 |
| TRDJ1 | Not detected – No cluster assigned | Fast | Cubic | 2.383 | .010 |
| NKG7 | NK-cells & T-cells - Adaptive immunity: Cytotoxicity | Fast | Cubic | 2.490 | < .001 |
| FCRL6 | NK-cells & T-cells - Adaptive immunity: Cytotoxicity | Fast | Cubic | 2.801 | < .001 |
| FGFBP2 | NK-cells & T-cells - Adaptive immunity: Cytotoxicity | Fast | Cubic | 2.929 | .002 |
| SULT1A1 | Neurons - Synaptic function & neurogenesis | Slow | Linear | 0.004 | .044 |
| CXCR2 | Neutrophils - Innate immunity signal integration | Slow | Linear | 0.004 | .017 |
| QPCT | Neutrophils - Phagocytosis & degranulation | Slow | Linear | 0.004 | .014 |
| AQP9 | Neutrophils - Chemotaxis & Apoptosis | Slow | Linear | 0.004 | .014 |
| FPR2 | Neutrophils - Chemotaxis & Apoptosis | Slow | Linear | 0.004 | .014 |

In the linear model, we found a total of 44 slow responding genes characterized by gradual changes over time following stress (**Fig. 2**, upper panels). Of these, 23 were upregulated and 21 were downregulated, with gene expression changes between the first and last measure varying between 0.60 to -0.48 counts per million per min (log_2_FC * 120 min). Using the cubic model, we found 14 fast responding genes characterized by non-linear trajectories, showing a significant peak immediately following stress before stabilizing at baseline levels at subsequent measurements (**Fig. 2**, lower panels). Four genes were found to be significant in both models. These genes initially peaked after stress, but declined below baseline at later measurements, leading to a negative linear slope being detected despite the overall non-linear trajectory.

**Fig. 2.**
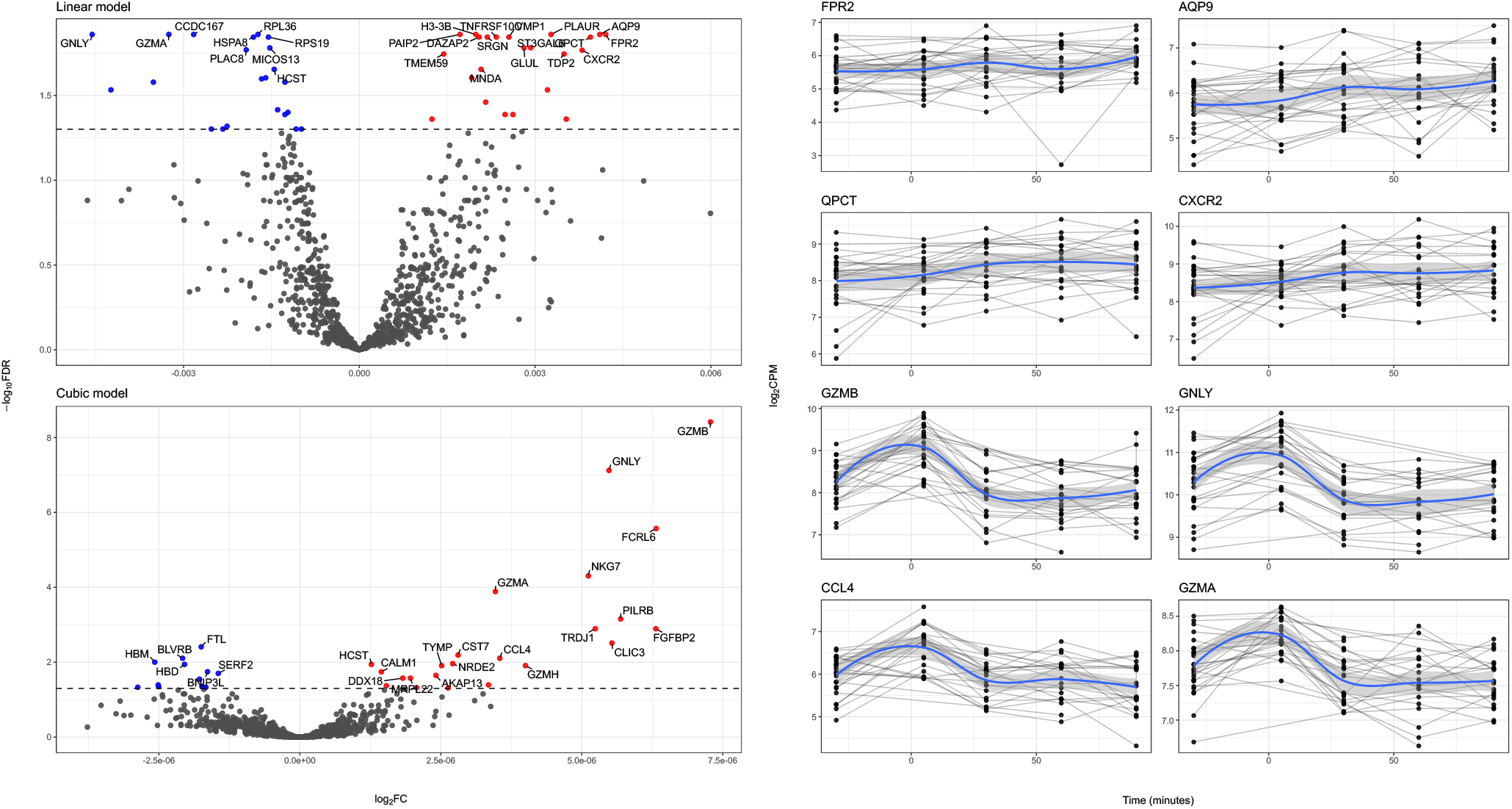
Time Dependent Gene Expression. Left panels: volcano plots of limma-voom differential expression models testing change over time. The upper plot shows the linear time model, and the lower shows the raw cubic time model. Points represent genes. The y-axis is -log_10_ FDR-corrected *p*-values, and the x-axis is log_2_ fold change. The dashed horizontal line indicates the false discovery rate threshold (FDR adjusted *p* < .05,). Highlighted and labelled genes denote the most significant hits. Right panels: expression trajectories for the top four genes from each model (linear: *FPR2, AQP9, QPCT, CXCR2*; cubic: *GZMB, GNLY, CCL4, GZMA*). Black points show voom-normalized expression (log2 transformed counts per million) at each sampling time with gray connecting repeated measures within individuals. The blue curve indicates loess-smoothed trend.

Overall, fast responding genes were associated with cytotoxic T cell and NK cell activity, and their gene expression trajectories mirrored that of lymphocyte cell counts (compare **Fig. 2** lower right panels with **Fig. 1C**). In contrasts, the slow responding genes were associated with neutrophil activity, but their trajectories did not follow those of neutrophil cell counts (compare **Fig. 2** upper right panels with **Fig. 1B**).

### Pathway, Transcription Factor and Interaction Analyses

Gene set enrichment analysis (GSEA) highlighted 20 enriched driver terms related to granzyme-mediated cytotoxicity, cytolytic granules and vesicles, and neutrophil degranulation (**Fig. 3**, full statistics available in **Supplementary Table S6**). Among the top ranked TFs identified by TFEA were multiple TFs related to IL-12 family signaling (e.g. *STAT4, TBX21, STAT5B, HLX* and *EOMES*. See **Supplementary Table S7** for complete list). Clustering of TFs and target genes separated two distinct clusters **(Fig. 4**): one cytotoxic T/NK-cell related cluster (including e.g., the genes *GZMA, GZMB, PLAUR, CCL4* and the TFs *STAT4, EOMES, PRDM1*) and one myeloid/neutrophil related cluster (including e.g., the genes *CXCR2, FPR2, AQP9, HRH2* and the TFs *STAT5B, NFIL3* and *ARNTL*). Protein-protein interaction analysis using STRING revealed significant enrichment above random expectation, indicating functional relatedness among the differently expressed genes (**Fig. 5**). Two separate interaction networks and enrichment were identified related to processes such as regulation of extrathymic T cell differentiation and NK cell mediated immunity, cytolytic granules, and cytotoxic T cell pyroptotic processes.

**Fig. 3.**
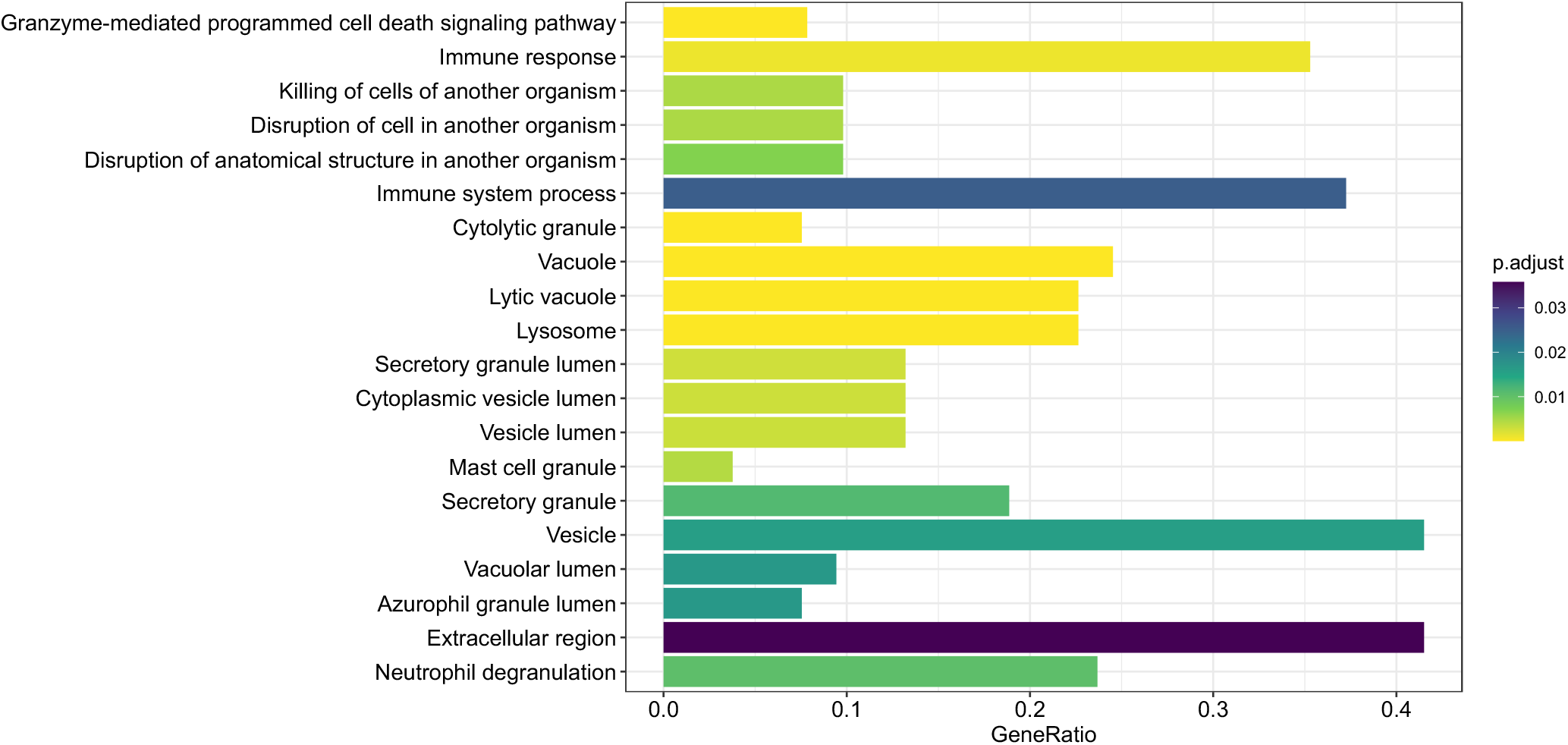
Enriched Terms and Pathways. Bar size on the x-axis indicates gene ratio (intersection size / query size). Color scale refers to significance levels of the gene set enrichment analysis results.

**Fig. 4.**
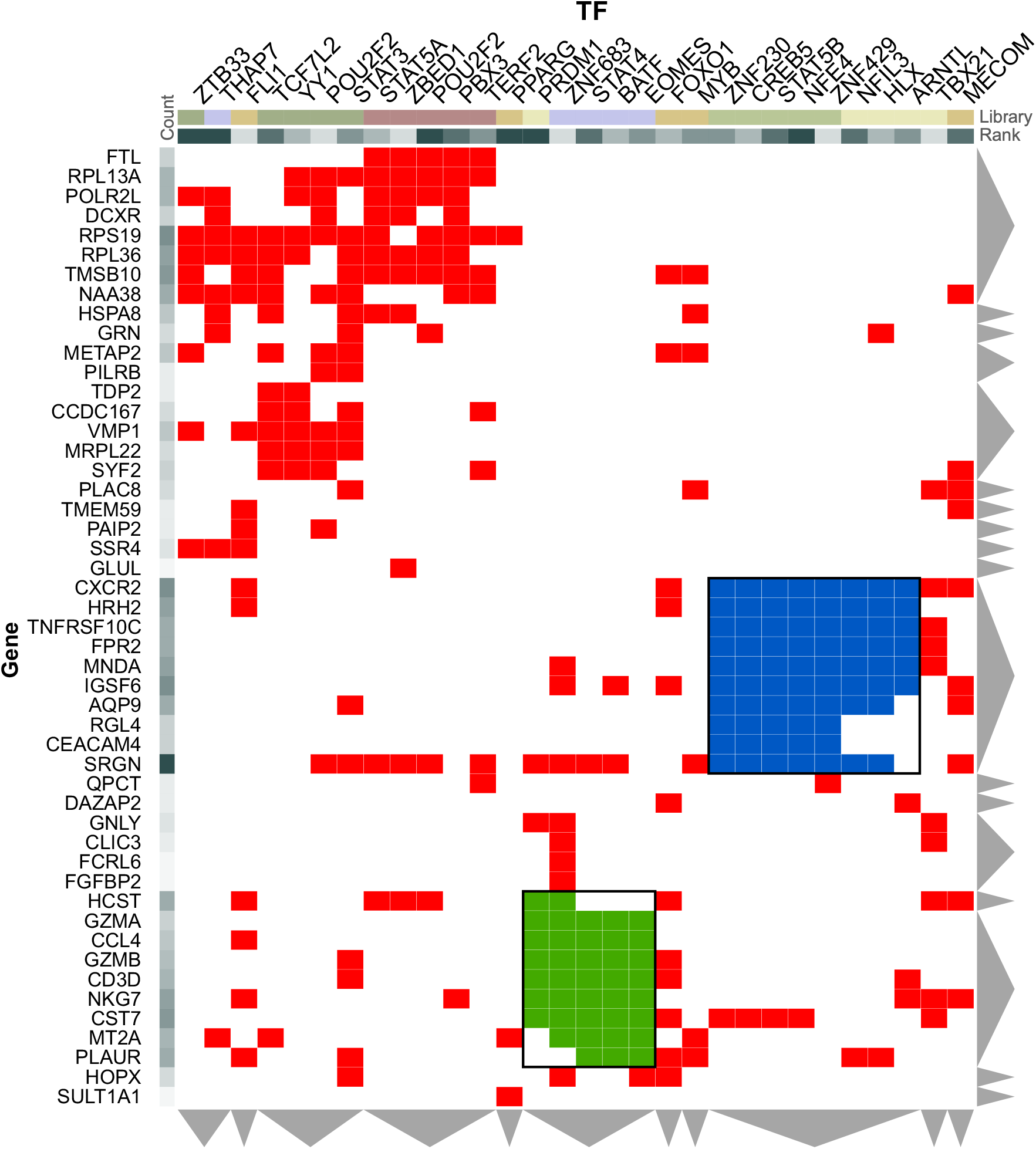
Transcription Factor and Target Gene Clustering. Differently expressed genes are shown on the x-axis and TFs on the y-axis. The cytotoxic T-/NK-cell related cluster has been highlighted blue, and the myeloid/neutrophil related cluster has been highlighted green.

**Fig. 5.**
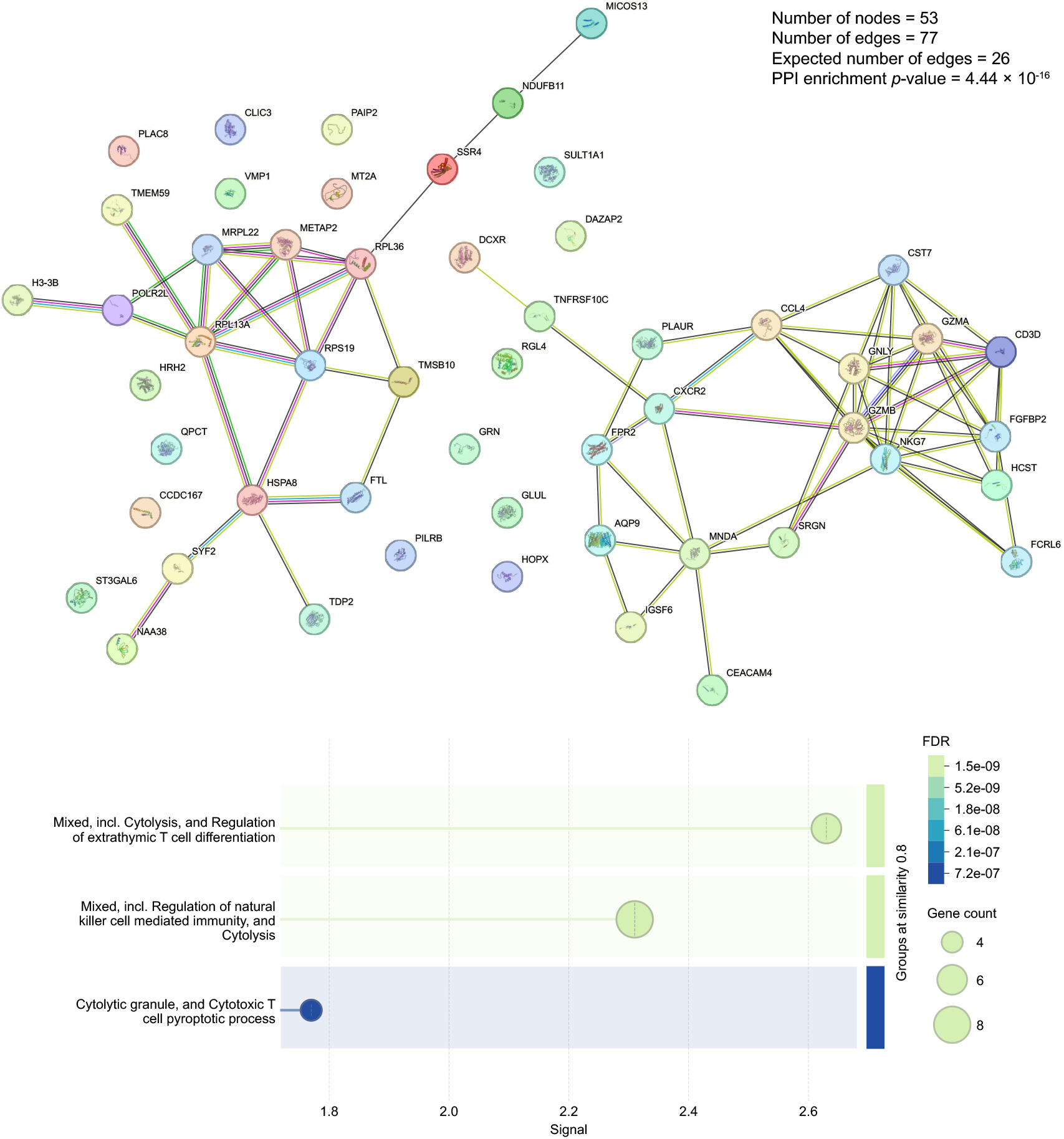
Protein-Protein Interactions. Upper: protein-protein interaction network constructed using STRING, showing known and predicted protein-protein interactions among nodes. Edges represent functional association. Lower panel: bubble plot summarizing gene ontology enrichment for STRING-defined local network clusters. Signal strength is shown on the x-axis, bubble size indicates gene count per term, and color corresponds to FDR adjusted *p*-values.

## Discussion

This study investigated rapid and delayed changes in gene expression following acute psychosocial stress in humans. We found that fast changes, with a distinct peak following stress, were related to cellular functions of NK-cells and T-cells, whereas slow and linear changes were related to neutrophil activity, consistent with recent mechanistic findings from mouse studies[14].

### Fast Gene Expression Responses to Stress

The rapid changes in gene expression, with a distinct peak immediately after stress, were mainly associated with NK and T cells, and parallelled the observed cell counts of lymphocytes (of which NK and T cells are subsets of). This suggests that, at least in part, the increased expression of these genes may be best explained by concurrent abundance of circulating lymphocytes. For example, we observed increases in the gene expression of granzymes (*GZMA, GZMB*), which are key components of granule-mediated cytotoxicity and expressed by multiple immune cells in peripheral blood, including cytotoxic T cells, NK cells, and dendritic cells[36, 37]. Granzymes are stored in cytolytic granules and can be released during immune activation[37, 38]. Consistent with this, we observed similar transient transcriptional patterns in *NKG7* and *GNLY*, which are both characteristic of cytotoxic lymphocytes and cytotoxic granule biology^37,39^. Thus, an increase in circulating cytotoxic lymphocytes could contribute to the observed transcriptional changes, even in the absence of strong cell-intrinsic upregulation.

### Slow Gene Expression Responses to Stress

We found a set of genes that showed gradual, linear upregulation with no pronounced peak following stress (e.g., *CXCR2, QPCT, AQP9, FPR2, SULT1A1* and *PLAUR*). Many of these genes are expressed predominantly in neutrophils and are involved in processes such as chemotaxis, migration, metabolism and degranulation. Notably, the trajectories of these genes did not consistently parallel the observed neutrophil counts. This dissociation suggests that the observed transcriptional changes cannot be fully explained by shifts in blood cell composition alone and instead point toward cell-intrinsic responses.

Activation of motor neurons during acute stress has been shown to increase neutrophil counts in mice 1 to 2 h later[14]. The activation of motor neurons released the *CXCR2* ligand *CXCL1* which is an attractor of neutrophils. We observed an increase in expression of *CXCR2*. We also observed an increase in the number of neutrophils in our sample 60 to 90 min following stress. The linear temporal pattern of *CXCR2* expression was not concordant with the biphasic trajectories of neutrophil counts, suggesting that *CXCR2* expression is cell intrinsic and not explained by cell counts alone. Recently, Dohi et al.[40] demonstrated that acute stress induced strong increases of *CXCL1/-2* and *CXCR2* in the cervical lymph nodes of male Sprague-Dawley rats which increased mRNA levels of *CXCR2*. The temporal pattern of *CXCR2* expression observed in our data showed a similar trend between baseline and post-stress timepoints. Based on these findings in rodents and our findings in humans, we speculate that IL-8 (human *CXCR2* ligand) release in blood following stress leads to increased neutrophil recruitment and succeeding upregulation of *CXCR2*.

### Biological Pathways and Transcription Regulators

Our combined analyses of protein interactions, gene sets, pathways and transcription factors suggest that immune activity in T-cells and neutrophils play a key role in the gene expression changes that follow acute stress. Our GSEA results revealed enrichment of biological processes related to cytotoxic and secretory immune functions, including granzyme-mediated programmed cell death, immune response, killing of cells of another organism, and several granule-associated gene ontology terms (cytolytic granule, secretory granule lumen, mast cell granula, and extracellular region). Similarly, our protein-protein interaction analysis indicated enrichment for processes such as regulation of extrathymic T-cell differentiation, NK-cell-mediated immunity, cytolytic granule formulation, and cytotoxic T-cell pyroptotic processes, all pointing to enhanced cytotoxic and inflammatory activity. Further analysis of inferred transcription factor regulation using ChEA3 highlighted IL-12 family-related regulatory signatures, suggesting that stress may transiently enhance Th1-type immune activation involving cytotoxic T-cells, NK-cells and neutrophils.

IL-12 is a key proinflammatory cytokine and a potent driver of type 1 immune responses, known for activating NK-cells[41, 42]. Beyond its immunological role, IL-12 contributes to neuroinflammatory processes implicated in major depressive and anxiety disorders[43]. On the other hand, it can either exacerbate or alleviate inflammation in central nervous system autoimmune conditions[41, 44]. In patients with panic disorder, IL-12 and IL-1β levels were found to correlate negatively with cortisol, indicating that stress-related hormonal responses may directly inhibit proinflammatory cytokine activity[45]. IL-12 primarily signals through IL-12-STAT4 activation, which has been shown to mediate most physiological effects of IL-12[46]. Consistent with these findings, our transcription factor analysis identified STAT4 as one of the top regulators. STAT4 was further linked to several genes associated with cytotoxic and inflammatory functions, including *GZMA, CCL4, GZMB, CD3D, NKG7, CST7, MT2A*, and *PLAUR*, indicating coherence between inferred transcriptional regulation and known IL-12/STAT4 immune programs.

Another notable transcription factor identified was *TBX21* (T-bet), a key regulator of lymphocyte differentiation and function that plays a pivotal role in programming CD8^+^-cells and NK-cells[47]. T-bet may also mediate behavioral responses to stress. In mice, deficiency of T-bet markedly reduced anxiety- and depression-like behaviors following acute stress exposure, accompanied by lower serum levels of the proinflammatory cytokines IL-6 and TNF-α^48^. This supports a potential genetic link, *TBX21* polymorphisms have also been associated with major depressive disorder[49].

### Clinical Implications for Anxiety and Depression

Cytokines have long been proposed to mediate the connection between stress, anxiety and depression[50]. The IL-12 family signaling pathway may be particularly important in this context due to its interaction with the HPA-axis[42]. Elevated plasma IL-12 levels have been reported in individuals with major depressive disorder[51], and treatment with antidepressants can reduce IL-12 concentrations in patients with depression[52] and anxiety^53^, although this reduction has not been observed in all studies[54]. Interestingly, drugs used to treat psoriasis that act on the IL-12 cytokine family, such as ustekinumab and guselkumab, have also been found to alleviate symptoms of anxiety and depression^48,55,56^. While these effects may partly reflect relief from psoriasis-related distress, they could also result from reduced IL-12 activity. Collectively, such findings suggest that pharmacological inhibition of the IL-12 pathway might hold potential as an adjunct therapy for depression and anxiety disorders[43, 57].

Our findings further identify specific genes that may help explain how stress relates to anxiety and depression, with particular attention to *PLAUR*. In a knock-out model, it has been shown that mice lacking uPAR (the protein encoded by *PLAUR*) display increased anxiety-like behaviors as compared to their wild type counterpart[58]. In humans, a recent UK Biobank study linked higher plasma *PLAUR* protein levels to an elevated risk of developing anxiety or depression over a 14-year period, and identifier *PLAUR* as a key inflammatory mediator in their comorbidity[59]. Elevated *PLAUR* levels have also been associated with current depression and PTSD in the same cohort^60,61^. In gene expression studies, *PLAUR* was among the core enrichment genes distinguishing men with generalized anxiety disorder from controls[62]. Consistently, increased *PLAUR* RNA expression has been observed in postmortem brains of individuals with major depressive disorder and PTSD[60], as well as across multiple neural and endothelial cell types in the dorsolateral prefrontal cortex[63]. Our results link these findings to the slower component of the stress response, which could suggest an involvement of neutrophils in anxiety and depression, as well as more general stress related processes occurring in many different cell types, including brain cells.

Another notable gene identified in our analyses was *GRN*, which encodes progranulin (PGRN), which has been implicated in multiple studies of anxiety. In mice, sex-specific differences in anxiety behaviors have been observed in wildtype, but not in Pgrn knockout animals, suggesting that PGRN contributes to the generation of sex differences in anxiety-related behaviors[64]. Progranulin also interacts with TNF-α receptors, and an imbalance between PGRN and TNF-α has been shown to induce memory impairments and anxiety-like behaviors in sleep-deprived mice^65^. Furthermore, treatment with either exogenous PGRN or a TNF-α inhibitor reversed these effects, while PGRN also suppressed dendritic spine loss and restored neurogenesis in a specific part of the hippocampus in these mice, indicating a neuroprotective role^65^. More recently, intracerebroventricular administration of PGRN following cerebral ischemia reduced anxiety-like behavior, and improved spatial learning, memory and hippocampal neurogenesis, suggesting its potential as a therapeutic candidate for mood and cognitive impairments after ischemic stroke[44]. Taken together, these findings implicate *GRN*/PGRN as a key factor in the interaction between stress, inflammation, and anxiety regulation.

We also observed gradual increased expression of glutamine synthetase (*GLUL*), which plays a key role in neurotransmission by recycling the excitatory neurotransmitter glutamate, and has been implicated in both stress and depression, as well as neurological disorders such as schizophrenia and epilepsy[66, 67]. During acute stress, glutamate release increases, followed by enhanced recycling through glutamine synthetase, a process tightly regulated by glucocorticoids[68]. Postmortem studies have shown decreased expression of *GLUL* in glial cells in individuals with depression, suggesting disruptions in glutamate metabolism may underly depressive pathology[69]. In male mice, lithium treatment showed sex-specific protection against stress-induced neurobiological changes, seen through a change in glutamine synthetase activity levels via a gene promoter reporter not observed in female mice[66]. Additionally, injection-induced stress alone increased promoter activity of *GLUL*, highlighting its sensitivity to acute stress exposure[66]. Altogether, these findings underscore the therapeutic potential of targeting the glutamate-glutamine cycle, as pharmacological modulation of glutamate synapses may offer alternatives conventional monoaminergic treatments for mood disorders[68, 70].

*FPR2* was another gene that increased slowly after stress. *FPR2* encodes the formyl peptide receptor 2 and is expressed by monocytes and neutrophils, where its expression is enriched during degranulation[71]. The FPR2 protein can recognize a wide variety of ligands, through which it can act as either a pro- or anti-inflammatory modulator and participate in the resolution of inflammation in the central nervous system[72]. In mice, *FPR* signaling has been shown to strongly impact anxiety-like behaviors^73,74^. For example, one recent mouse study demonstrated that treatment with an *FPR* antagonist (synthetic peptide WRW4) mitigated anxiety- and depression-like behaviors in a social isolation mouse model^74^.

Several of the genes that changed expression rapidly after stress have also been linked to anxiety and depression. For example, downregulated *GZMB* expression has been observed in individuals with current depression as compared to those in remission[75], while blood levels of the cytokine CCL4 has been found to be elevated in individuals with anxiety, depressive disorders, and *PTSD*[76–78]. This suggests that both genes that change expression shortly after stress, and genes that show more delayed expression changes, have been found to be associated with stress-related disorders.

In summary, our results show that acute social stress induces both rapid and delayed transcriptional changes across immune pathways. The early response reflects activation of cytotoxic lymphocyte programs, while the later response involves neutrophil-associated transcriptional regulation, both implicating IL-12 family signaling associated regulatory programs. These findings reveal a coordinated temporal structure of immune gene activation following psychosocial stress and identify molecular targets such as STAT4 and *PLAUR* that may bridge peripheral immune activation with neurobiological processes underlying anxiety and depression.

## Acknowledgements

This work was supported by the Swedish Research Council (grants no. 2018-01322 and 2023-01093) and the Bank of Sweden Tercentary Foundation (grant no. P25-0219).

