## Supplemental Figures for "Fast and Slow Gene Expression Changes in Blood Following Acute Social Stress"

### Supplementary Figure S1 STRT Outlier Detection

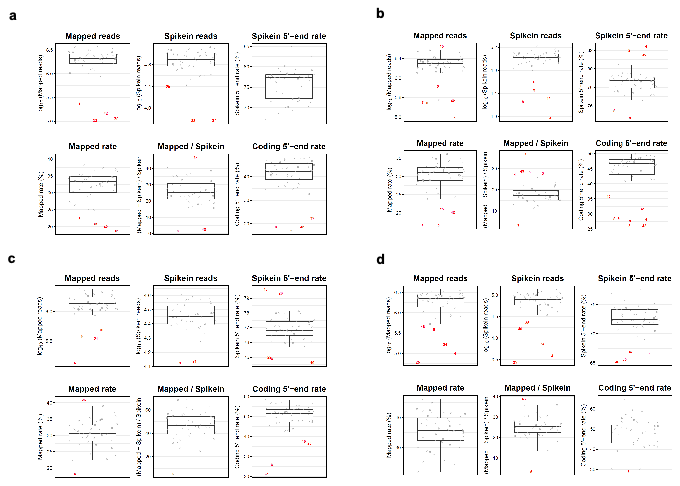

*Detection and exclusion of outlier samples across four STRT-seq libraries.* *Each panel (a–d) represents one sequencing library (Library 1 - Library 4) and shows quality control (QC) metrics: mapped reads (log10), spike-in reads (log10), spike-in 5′-end rate, mapped rate, mapped/spike-in ratio, and coding 5′-end rate. Red dots indicate samples flagged as outliers based on unusually low or high values in one or more QC metrics.*

*Outlier samples were excluded from downstream analyses:*

*(a) Library 1: samples 8, 25, 33, 34, 37 and 42*

*(b) Library 2: samples 1, 2, 3, 4, 5, 7, 8, 9, 14, 15, 40, 42 and 43*

*(c) Library 3: samples 1, 6, 10, 19, 20, 21, 22, 23, 25, 26 and 40*

*(d) Library 4: samples 4, 5, 25, 34, 35, 43, 46 and 47.*

### Supplementary Figure S2 Library Bias Assessment

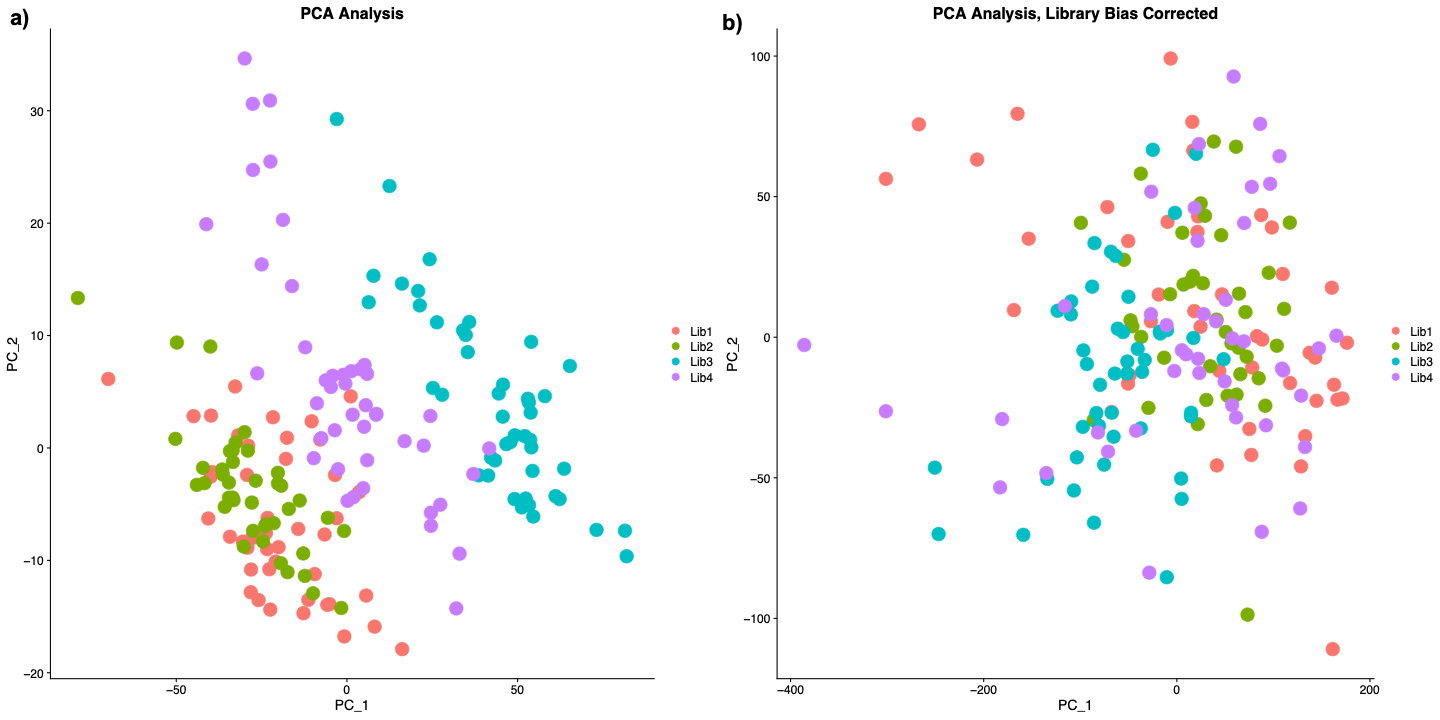

*Library bias assessment before and after correction. Left panel: principal component analysis (PCA) shows clear clustering of samples by library before correction, indicating strong library-associated bias. Right panel: after library bias correction, the samples no longer cluster by library, suggesting that the batch effect has been removed. Therefore, library information was not included as a covariate in the final differential gene expression model.*
